# Evolutionary causes of lifespan extension by dietary restriction: linking theory and mechanisms

**DOI:** 10.1101/2020.01.14.904599

**Authors:** Laura M. Travers, Hanne Carlsson, Elizabeth M. L. Duxbury, Alexei A. Maklakov

## Abstract

Dietary restriction (DR), reduced food intake without malnutrition, increases lifespan across a broad range of taxa, but the evolutionary underpinning of this phenomenon is poorly understood. The resource reallocation hypothesis proposes that dietary restricted animals divert resources from reproduction to somatic maintenance to increase survival in times of nutrient scarcity in favour of future reproduction. The “longevity by-product” hypothesis proposes instead that dietary restricted animals increase nutrient recycling via autophagy to maximise immediate reproduction, thereby reducing cellular toxic waste and leading to longer lifespan as an unselected by-product. The “longevity by-product” hypothesis makes a unique prediction that blocking autophagy in DR animals will simultaneously reduce lifespan and reproduction. To test the adaptive value of autophagy under dietary restriction, we inhibited autophagy using *bec-1* RNAi knockdown in DR and fully-fed *Caenorhabditis elegans* nematodes. Our findings confirm that autophagic inhibition results in a significantly shorter lifespan under DR, suggesting that autophagy is important for survival in times of famine. Remarkably, we also show that inhibiting autophagy throughout adult life significantly increases reproduction in both dietary restricted and fully fed worms. Moreover, this did not come at a transgenerational cost to offspring fitness. Our results suggest that autophagy is an energetically costly process that reduces resources available for reproduction, but is necessary for survival during famine, and are thus consistent with the resource reallocation hypothesis.

## Introduction

Dietary restriction (DR), a moderate reduction in food intake without malnutrition, extends lifespan in many organisms, from yeast to primates ^1–4^. DR is the most reproducible environmental intervention to increase lifespan and improve health outcomes in a wide range of species ^5–7^. Research on genes that modulate lifespan has revealed that nutrient-sensing pathways play a key role in the physiological changes that affect ageing and lifespan in response to nutrient availability ^8–10^. Changes in activation of nutrient-sensing pathways in response to nutrient availability are thought to reflect an adaptive phenotypically plastic response, allowing organisms to optimise investment between somatic maintenance and reproduction to maximise fitness in varying environments ^11,12^ The predominant evolutionary theory, the resource reallocation hypothesis, proposes that the physiological effect of DR, characterised by increased lifespan and reduced reproduction is an adaptive strategy to increase survival in times of resource scarcity ^12^ By diverting resources away from reproduction and towards somatic maintenance, organisms can maximise their chances of survival by reducing investment into energetically costly activities such as reproduction, and prioritise getting through temporary famine by investing in somatic maintenance. According to this theory, when nutrients become plentiful again, organisms can reverse the allocation from somatic maintenance, back into reproduction ^12^.

However, several shortcomings of the resource reallocation theory have recently been highlighted. While DR commonly results in a decrease in reproduction, recent studies have shown that reduced reproduction is not a requisite for increased lifespan (for a review, see ^13,14^). For example, DR can extend lifespan in sterile and non-reproducing animals ^3,15–17^ which suggests that lifespan extension may occur in the absence of resource allocation from reproduction to somatic maintenance. Experimental evolution in *D. melanogaster* has also found no evidence for coevolution between longevity and reproduction under DR ^18,19^ These findings highlight that extended lifespan under DR may not be constrained by direct competition for limited resources as suggested by the resource reallocation theory.

In addition to the evidence disputing the mechanistic underpinnings of resource reallocation, Adler & Bonduriansky ^20^ challenge the evolutionary explanation of the adaptive resource reallocation theory, that is, that the DR response functions to increase survival over reproduction. One of their key criticisms is that increasing investment into somatic maintenance to promote survival, while significantly decreasing or “shutting down” reproduction, is not adaptive. Given that selection should favour reproduction over longevity, they argue that delaying reproduction during food shortages would not be adaptive in the wild where extrinsic mortality is high for many species ^20^. In summary, the evidence above casts doubt on whether dietary restricted animals strategically allocate resources between somatic maintenance and reproduction to improve survival in times of famine.

Adler & Bonduriansky ^20^ propose an alternative evolutionary explanation for the DR response by focusing on the physiological cell protection and repair mechanisms that are triggered in response to DR. One key cellular process involved in removing and recycling damaged cell components that is upregulated in response to DR is autophagy. Autophagy is a process whereby portions of the cytoplasm, including misfolded proteins, mitochondria, and other organelles, are degraded, catabolized and recycled ^21^. Large double-membrane vesicles called autophagosomes are generated, which engulf cytoplasmic organelles. The autophagosomes fuse with lysosomes where the breakdown products are recycled to the cytoplasm. Adler & Bonduriansky ^20^ argue that the primary evolutionary function of increased autophagy under DR is not to improve somatic maintenance to maximise survival under starvation, but instead to free up nutrients through increased nutrient recycling to invest in reproduction. If delaying reproduction is not adaptive due to the risk of extrinsic mortality, upregulating autophagy would allow the organism to make the most efficient use of scarce resources for some immediate reproduction. In this scenario, extended lifespan in response to DR is merely a non-adaptive by-product of increased autophagy. According to this theory, increased autophagy is beneficial by making more efficient use of resources when nutrients are scarce, which results in the availability of more nutrients for reproduction that would otherwise not be available ^20^.

The evolutionary question of why selection has favoured the up- and downregulation of autophagy in response to nutrient availability remains unanswered. We aimed to test whether the primary role of increased autophagy, one of the key mechanisms involved in extending lifespan under DR ^22–28^, is to increase survival as suggested by the resource reallocation theory ^12^, or to increase immediate reproduction as proposed by the longevity by-product hypothesis ^20^. The longevity by-product hypothesis posits that increased autophagy frees up resources for reproduction. Therefore, this theory makes the unique prediction that reproduction should decrease if autophagy is inhibited in DR conditions because the extra nutrients normally made available through upregulated autophagy would not be available to invest in reproduction here. In this study, we tested the adaptive significance of increased autophagy under DR. We used RNA interference (RNAi) to inhibit a key autophagy gene in dietary restricted and ad lib fed *Caenorhabditis elegans* worms, and examined the consequences for reproduction and lifespan. We also tested for potential transgenerational consequences of inhibiting autophagy in fully fed worms by measuring reproduction and lifespan in the offspring of ad lib fed parents.

## Methods

### Strains

We used Bristol N2 wild type and *eat-2 (ad 1116)* mutant (genetic model of DR) strains of *C. elegans* nematodes in all assays. *Eat-2* mutants have a defect in a nicotinic acetylcholine receptor subunit that functions in the pharyngeal muscle ^29^, resulting in reduced pharyngeal pumping rates and 20% less food intake ^30^. Due to dietary restriction, *eat-2* mutants live approximately 20% longer and have fewer offspring (~50%) than wild type worms ^15,27^.

Before the start of the experiment, we bleached plates to collect eggs from worms recovered from frozen. We used standard NGM agar plates to grow the nematode populations ^31^, with added antibiotics (100 μg/ml Ampicillin and 100 μg/ml Streptomycin) and a fungicide (10 μg/ml Nystatin) to avoid infections ^32^ Before the experiment began, we fed the nematode populations antibiotic resistant *Escherichia coli* OP50-1 (pUC4K), gifted by J. Ewbank at the Centre d’Immunologie de Marseille-Luminy, France. From defrosting and throughout the experiment, we kept worms in 20°C and 60% relative humidity climate chambers. We used RNAi to knock down Beclin-1, an ortholog of mammalian Beclin1, a gene that is part of an operon that contains the stress-inducible transcription factor gene *skn-1* which is essential for the lifespan extension caused by DR ^33^. RNAi *bec-1* was provided by the Julie Ahringer RNAi library. Previous studies have shown inactivation of *bec-1* in *eat-2* mutants abrogates the lifespan extension effects of DR ^22,26,27^. Thus, BEC-1 is required for the lifespan extension induced by dietary restriction in *eat-2*. While inhibiting *bec-1* prevents food limitation from extending lifespan, it has also been shown to have little effect on the lifespan of well-fed animals ^27^.

In order to induce RNAi knockdown, we fed the nematodes strain HT115 (DE3) of *E. coli* containing a Timmons and Fire feeding vector L4440 modified to express the dsRNA of *bec-1.* In addition to the RNAi knockdown bacteria, we used a strain of HT115 carrying an empty L4440 feeding vector to feed to worms in our control treatment ^34^. We grew cultures of the RNAi clones in LB medium supplemented with Ampicillin (50 μg/ml) and then seeded onto 35mm standard NGM plates with the addition of IPTG (1 mM) and Ampicillin (50 μg/ml), as recommended by Kamath et al. ^35^. After seeding plates with either RNAi *bec-1* knockdown or the control empty vector RNAi, we grew the bacteria overnight in 20°C to induce expression of RNAi, before placing worms on the plates.

### Experimental design and set-up

For logistical reasons, we conducted the lifespan assay in three experimental blocks and the reproduction assay in four blocks. We collected age-synchronized eggs from N2 and eat-2 worms at peak reproduction (Day 2 in N2 and Day 3 in *eat-2*) from unmated (i.e. self-fertilised) hermaphrodites fed on OP50-1 (pUC4K). Eggs from each strain were placed on OP50-1 plates to develop from egg to late L4/Day 1 adults. We placed eggs onto the development plates, which we assigned a random identifier (e.g. 1-10) to blind the experimenter to the worm strain (N2 and *eat-2*) henceforth.

At the beginning of the experiment, age-synchronised (Day 1) adult worms from both strains were randomly selected from the development plates and placed on either RNAi *bec-1* seeded plates or on empty vector control plates and maintained on their respective plate type throughout life. We used a combination of randomisation and stratification to set-up and handle experimental plates throughout the experiment, with plates assigned random identifiers (e.g. 1–100). Thus, experimenters were blind to worm treatment and strain during the daily transfers when we scored lifespan, and during offspring counting.

### Offspring production

To obtain data on the number of offspring produced by each worm, we reared single unmated (i.e. self-fertilised) hermaphrodites on individual plates from adult Day 1 until reproduction ceased. We moved the worms onto new plates every 24h, incubated eggs for 48hrs at 20°C, and counted the number of progeny. For each block, 15–30 worms were set up for each treatment-strain combination. Over the four blocks, we assayed a total of 69–85 worms per treatment-strain combination.

### Lifespan

Lifespan assays were performed on unmated hermaphrodites from adult Day 1 until death. We set up worms in groups of ten and transferred onto fresh plates daily while scoring survival. Death was defined as the absence of movement in response to touch. For each block, we set up 50–100 worms for each treatment-strain combination. Over the three blocks, we assayed a total of 170 worms per treatment-strain combination.

### Offspring reproduction and lifespan assays

To explore the transgenerational consequences of inhibiting autophagy in parents, we measured progeny production and lifespan in the offspring of fully fed N2 worms exposed to RNAi *bec-1* in adulthood. We collected age-synchronized eggs from RNAi *bec-1* and control RNAi empty vector fed N2 worms at peak reproduction (Day 2). At the beginning of both the offspring lifespan and reproduction assay, age-synchronised (Day 1) adult worms were haphazardly selected from the development plates and placed on standard NGM agar plates seeded with 0.1ml of *Escherichia coli* OP50-1 (pUC4K) and maintained on OP50-1 plates throughout the duration of both assays. In the reproduction assay, we placed single unmated (i.e. self-fertilised) hermaphrodites on individual plates from adult Day 1 until reproduction ceased. For the lifespan assay, we set up worms in groups of ten and transferred onto fresh plates daily while scoring survival. Both assays were conducted in two experimental blocks. As with the parental assays, offspring assays were conducted randomised and blinded, as described above.

### Statistical analyses

We used R version 3.5.1 ^36^ for all statistical analyses and figures. We analysed lifetime reproductive success (LRS; total number of offspring produced per individual) in both the parents and offspring assays using linear mixed effects models (LMM) implemented in the *lme4* package version 1.1-21 ^37^ For parental LRS, we fitted treatment and strain, and their interaction as fixed effects, along with the development plate (plate on which the individual worm developed from egg to adult prior to the beginning of the experiment) nested within experimental block as random intercepts. For offspring LRS, we fitted treatment and block (centred) as fixed effects, as well as the development plate nested within experimental block as random intercepts. We obtained P values from LMMs from t-tests using Satterthwaite’s approximation for denominator degrees of freedom implemented in *lmerTest* ^38^. We also analysed LRS by bootstrapping using the *dabestr* package ^39^. The 95% confidence intervals are graphically represented by the package in Figures 1(a) (parents) and 3(a) (offspring).

**Figure 1 (a):**
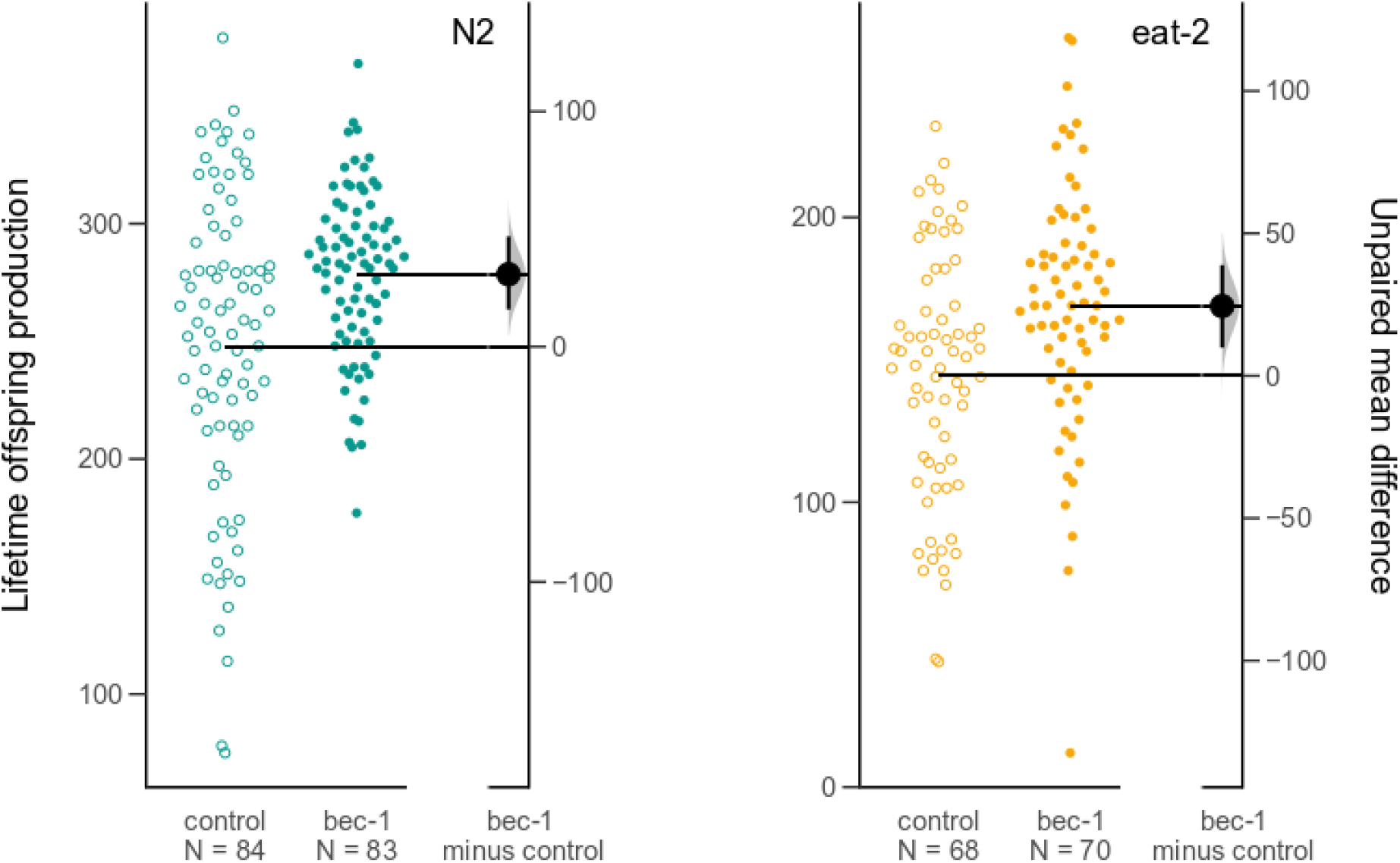
Inhibition of autophagy increases reproduction in N2 and eat-2 worms. Estimation plots of total lifetime reproduction for N2 (green) and eat-2 (yellow) where RNAi bec-1 treatment (filled circles) are compared to the control (open circles), with a graded sampling distribution of bootstrapped values and bootstrapped 95% CI, implemented in dabestr ^39^.

**Figure 1 (b):**
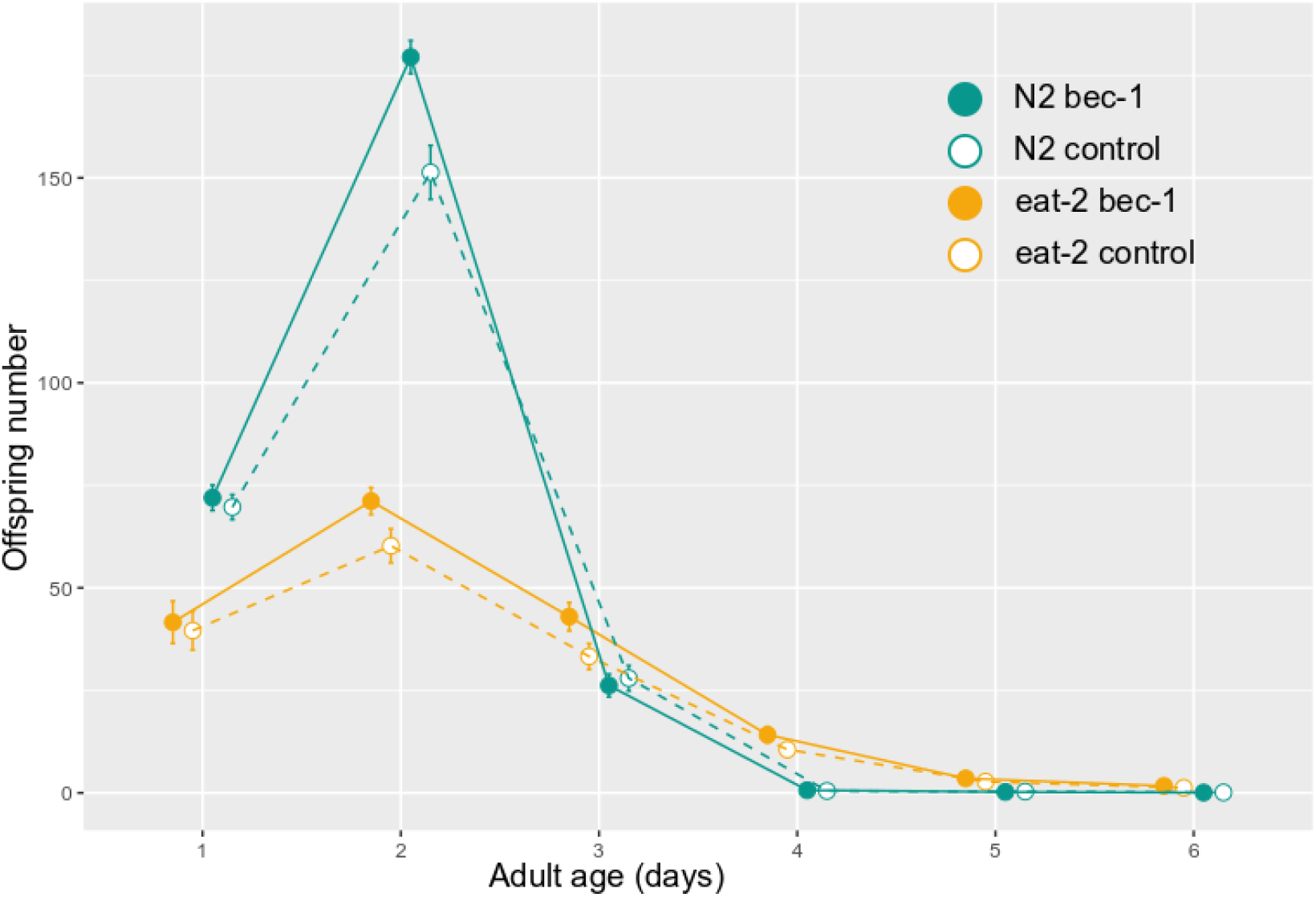
Age-specific reproduction. Age-specific reproduction for each nematode strain and RNAi treatment combination; circles and error bars represent mean ± SE’s. Colours indicate N2 (green) and eat-2 (yellow) worm strains. Open circles and dashed lines depict offspring number from control empty vector treatment, filled circles and solid lines depict RNAi bec-1 treatment.

We analysed lifespan in both parents and offspring using mixed effects Cox proportional hazard models implemented in the *coxme* package version 2.2-14 ^40^. Individuals that died of matricide (internal hatching of eggs) or escaped from agar were censored. For parental lifespan we fitted treatment and strain (both centred), and their interaction as fixed effects. We centred the treatment and strain by coding factor levels as minus 0.5 and 0.5, respectively, to facilitate the interpretation of main effects in the presence of an interaction ^41^. We included random intercepts for experimental blocks and group plate to account for variation due to shared environments. For offspring lifespan, we fitted treatment and block (centered) as fixed effects (offspring were all from the N2 strain) along with group plate as a random intercept.

## Results

We found a significant effect of treatment (RNAi bec-1 and empty vector control) on lifetime reproduction in both the *eat-2* dietary restricted mutants and N2 wild type ad lib fed worms. *Eat-2* and N2 worms that were fed RNAi *bec-1* produced on average 19% and 11% more offspring, respectively, compared to control worms fed the empty vector (t 1, 250.5 = 5.69, p < 0.001: Figure 1 and Table 1). There was no significant interaction between treatment and nematode (N2 and *eat-2*) strain (t_1, 248.9_ = −0.83, p = 0.406), hence we removed the interaction from the final model. As expected, *eat-2* worms produced fewer offspring (mean = 144 ± 5.30 SE) than the wild type N2 (mean = 248 ± 7.04 SE; t _1, 61.9_ = −17.14, p < 0.001).

**Table 1:** Full model summary for lifetime reproduction. Coefficients, standard errors, test statistics and variance components are taken from a LMM on the number of offspring produced. Effects associated with a p-value smaller than 0.05 are highlighted in bold.

| Lifetime reproduction ($N = 305$) | | | linear mixed effect model | | |
| --- | --- | --- | --- | --- | --- |
| Fixed effects | Coef | SE<br>(coef) | ddf | t | p |
| Intercept | **252.607** | **11.628** | **3.783** | **21.724** | **<0.001** |
| treatment | **-106.26** | **6.2** | **250.536** | **5.687** | **<0.001** |
| strain | **27.746** | **4.8** | **61.987** | **-17.14** | **<0.001** |
| Random effects | Var | SD |  |  |  |
| Block (4 levels) | 441 | 21 |  |  |  |
| Development plate | 232.9 | 15.26 |  |  |  |
| Residual | 1782.1 | 42.21 |  |  |  |

We found a marginally non-significant interaction between treatment and strain for lifespan (z = 1.94, df = 1, p < 0.053). Inhibition of autophagy reduced lifespan in DR worms substantially more than in N2 worms (proportional hazard odds ratios 2.06 versus 1.3). This means DR worms had a two-fold increase in mortality risk when we inhibited autophagy, while autophagy inhibition increased mortality in N2 worms only by a third (Figure 2 and Table S1).

**Figure 2:**
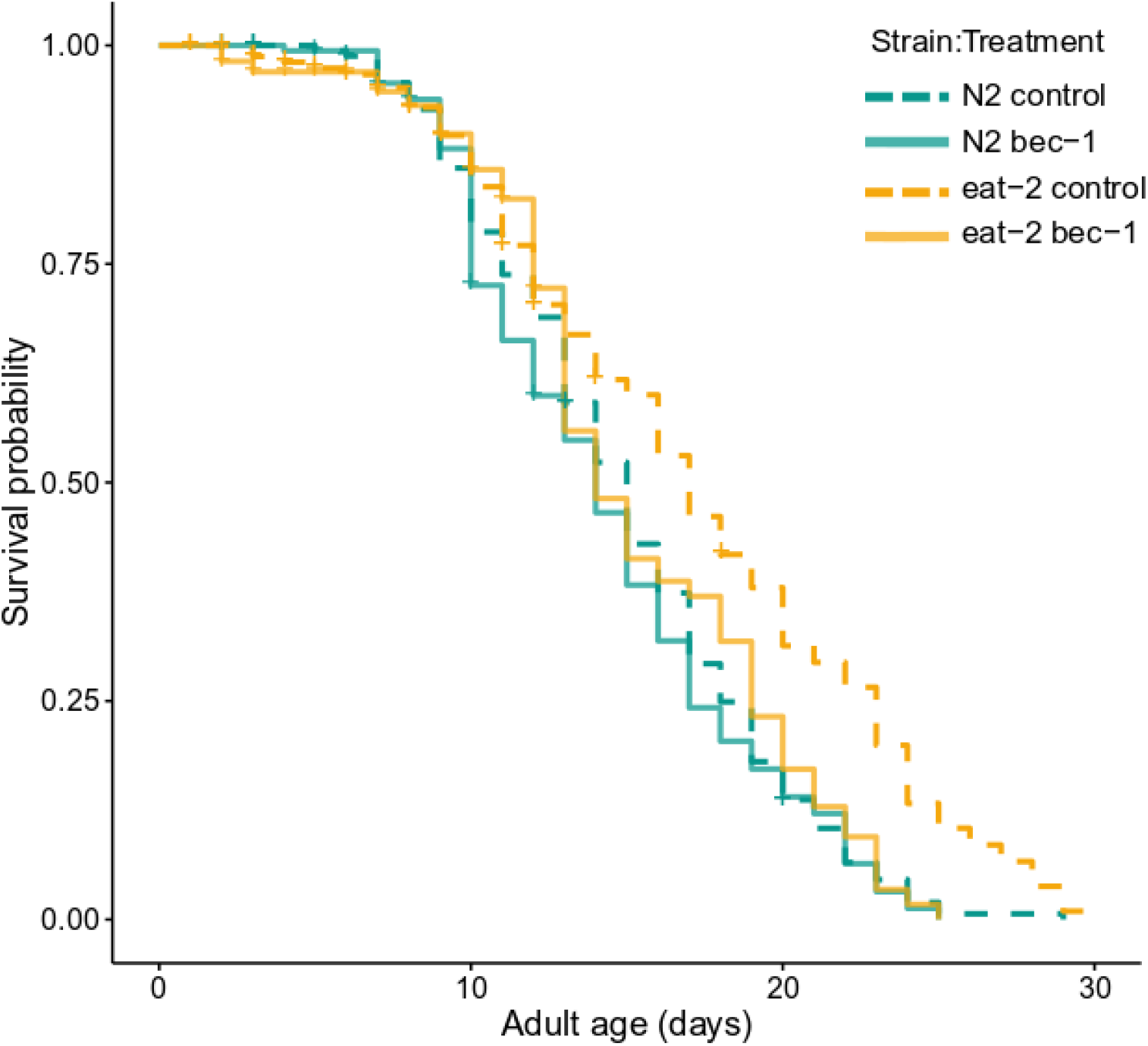
Reduced survival in eat-2 worms when autophagy is inhibited. Survival curves for each nematode strain and RNAi treatment combination. Colours indicate N2 (green) and eat-2 (yellow) worm strains. Solid lines depict RNAi bec-1 treatment, dashed lines depict control empty vector treatment. Plus-signs indicate censored individuals (matricide or escape; see main text).

We found no effect of parental inhibition of autophagy on either LRS (t _1, 93.4_ = −0.94, p = 0.349: Figure 3 (a) and Table S2) or lifespan (z= −0.50, df = 1, p = 0.620: Figure 3 (b) and Table S3) in the offspring of fully fed N2 worms.

**Figure 3(a):**
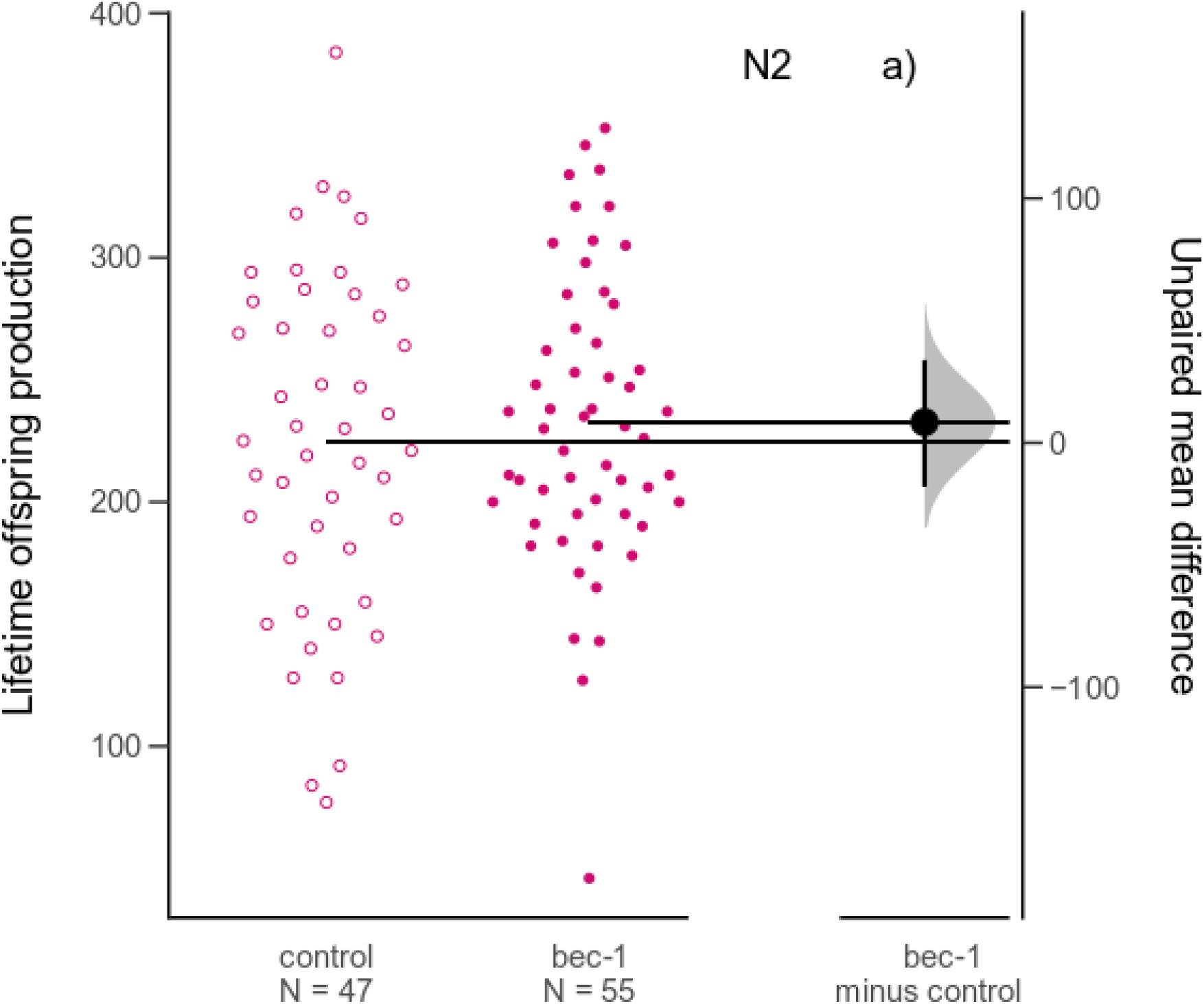
No difference in reproduction in offspring of N2 parents when autophagy is inhibited compared to controls. Estimation plots of total lifetime reproduction for offspring of N2 parents. Offspring from RNAi bec-1 treated N2 worms (filled circles) are compared to offspring from the control empty vector treated N2 worms (open circles), with a graded sampling distribution of bootstrapped values and bootstrapped 95% CI, implemented in dabestr^39^.

**Figure 3 (b):**
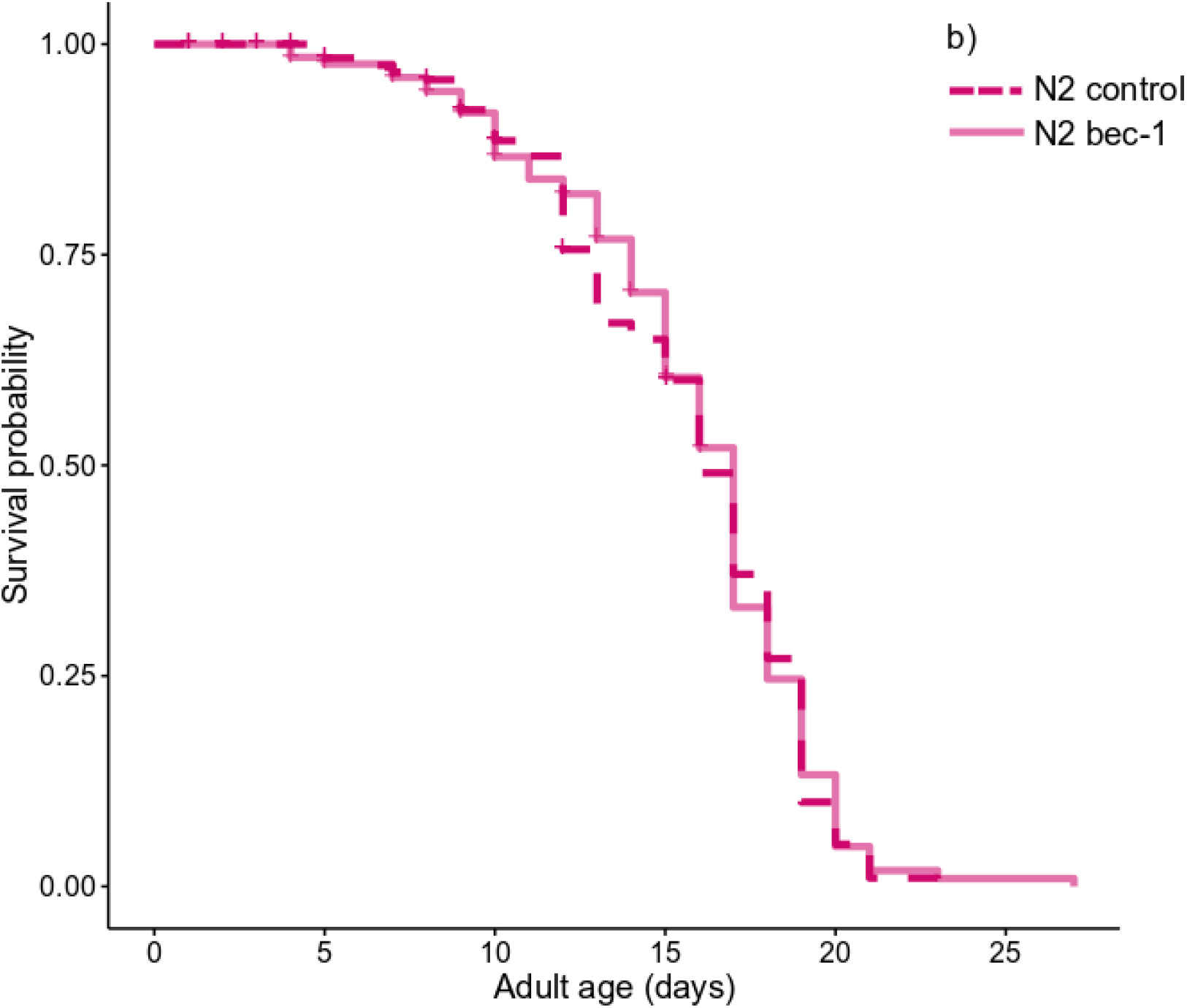
No difference in lifespan in offspring of N2 parents when autophagy is inhibited compared to controls. Survival curves for N2 offspring. Solid lines depict offspring from RNAi bec-1 treated N2 parents, dashed lines depict offspring from control empty vector treated N2 worms. Plus-signs indicate censored individuals (matricide or escape; see main text).

## Discussion

We tested the adaptive significance of increased autophagy in response to nutrient scarcity. For many years, it had been assumed that autophagy acts to promote survival by maintaining cellular homeostasis during periods of famine ^42^ A more recent suggestion is that autophagy may be activated during famine to free up resources for immediate reproduction ^20^. A unique prediction of this “longevity by-product” hypothesis is that blocking autophagy under DR should reduce reproduction compared to DR controls with intact autophagy. We inhibited the autophagy gene *bec-1*, previously shown to be required for lifespan extension under DR ^22–27^, and examined the consequences for reproduction and lifespan in DR and fully fed worms. Contrary to the prediction of the “longevity by-product” hypothesis, we found a substantial increase of 10–20% in offspring number when autophagy was inhibited, both in DR and in fully fed worms. Moreover, we confirmed previous findings that RNAi *bec-1* significantly shortens lifespan in *eat-2* (DR) mutants but has a lesser effect on lifespan in N2 wild type worms ^26,27^. Our findings support the adaptive explanation proposed by the resource reallocation hypothesis, which posits that increased autophagy under DR is adaptive to increase survival during famine.

In this study, we aimed to test why selection has favoured the up- and downregulation of autophagy in response to nutrient availability. While our findings support the evolutionary explanation proposed by the resource reallocation theory, the mechanistic underpinnings of lifespan extension under DR remain unclear. Recent research suggests that reproduction and lifespan are not regulated through direct competition for resources, but are mediated through molecular signals ^1,14,43–46^. When an animal is fully fed, the activation of nutrient-sensing pathways inhibits cellular recycling and repair mechanisms. Inhibited autophagy allows cell proliferation and growth, allowing an organism to increase reproduction ^1^. Nutrient-sensing pathways are deactivated in response to nutrient scarcity, which results in the upregulation of cellular recycling mechanisms ^2^. Thus, while increased autophagy extends lifespan through repair of damaged cells, it also limits cellular growth and proliferation ^47,48^, which limits reproduction. We found that inhibition of autophagy resulted in an increase in reproduction in both DR and fully fed worms, suggesting that increased cell growth and proliferation resulting from decreased autophagy in both DR and wild type worms allows individuals to increase their reproduction. Thus, increased autophagy under nutrient scarcity does not serve to increase reproduction, instead showing the opposite effect. Autophagy may use up more energy than it produces, which could explain why we found an increase in reproduction when autophagy was suppressed in both DR and wild type worms.

Inhibition of other autophagy genes also shortens lifespan in DR and nutrient-sensing-deficient animals. Similar to the effects of *bec-1* inhibition, suppression of *atg7* reduces lifespan in *eat-2* mutants, but has little effect on the lifespan of N2 wild type worms ^26^. Inhibition of *bec-1*, *atg-7* and *atg-12* reduces longevity in long-lived *daf-2 C. elegans* mutants ^24,25^. Toth et al. ^22^ also found that mutants with reduced insulin/IGF-1 signalling and dietary-restricted worms require genes involved in autophagy to live long. Collectively, these findings suggest that autophagy plays a key role in regulating lifespan in response to nutrients.

A key question emerging from our findings is why basal levels of autophagy are higher than what seems optimal for maximising reproduction in fully fed control worms. Increased reproduction when autophagy is inhibited in wild type worms suggests that lower expression of *bec-1* should be beneficial for offspring production while incurring no costs to survival when worms are fully fed. So why do worms not plastically decrease the rate of autophagy (expression of *bec-1*) during times of plenty to increase their reproductive output? The important role of BEC-1 in various biological processes may shed light on why expression is not downregulated to maximise reproduction in full feeding. In *C. elegans*, BEC-1 is required for embryo development and viability, dauer formation and germline formation ^25,49–53^. Given the important role of BEC-1 during development, the lack of downregulation may be an example of sub-optimal gene expression in adulthood due to the diminishing strength of selection with age ^54–56^.

BEC-1 is also important for DNA repair; *bec-1* null and *bec-1* RNAi fed animals show increased accumulation of DNA damage and an increase in germline cell death due to a delay in apoptotic cell corpse degradation ^49,57^ *C. elegans* with defects in DNA damage repair display a reduction in the number of viable eggs laid after DNA damage^58^. Therefore, while downregulation of autophagy increases the number of offspring produced, it could result in decreased offspring quality due to diminished ability to maintain the germline. However, our investigation of lifetime reproduction and lifespan in the offspring of fully fed wild type *bec-1* fed parents did not reveal any transgenerational costs of parental autophagy inhibition.

## Conclusions

In summary, our findings suggest that autophagy is increased in response to nutrient scarcity to improve survival rather than to maximise reproduction. Therefore, they support the resource reallocation theory for why DR extends lifespan, and refute the idea that autophagy frees up resources to maximise reproduction during famine. Increased autophagy activated by decreased insulin signalling limits cell proliferation, which could trade-off with cell repair and processes that require cell proliferation such as reproduction.

Our findings suggest that reproduction could be increased with little cost to survival if autophagy is downregulated during full feeding in adulthood. The fact that fully fed wild type worms do not decrease autophagy despite the apparent potential for increasing reproductive fitness points towards hidden costs of autophagy inhibition. Future studies could further explore the impact of inhibiting parental autophagy on offspring quality by measuring the fitness consequences in a variety of stressful environments.

## Author contributions

LMT and AAM conceived and designed the study. LMT, HC, and EMLD collected the data. LMT analysed the data and wrote the first draft of the manuscript. AAM revised and edited the manuscript, and all authors approved the final version.

## Acknowledgements

Tom van Baalen and Kris Sales assisted with laboratory work. We thank Andreas Sutter for advice on statistical analyses. This work has been supported by ERC Consolidator Grant GermlineAgeingSoma to AAM.

