## Supplementary material for "Evolutionary causes of lifespan extension by dietary restriction: linking theory and mechanisms": Table S3

**S3:** Full model summary for lifespan of N2 offspring treated with RNAi *bec-1* versus controls.

| **Lifespan in N2 offspring** ***(N = 212)*** | | **coxme** | | |
|---|---|---|---|---|
| **Fixed effects** | **Coef** | **SE (coef)** | **z** | **p** |
| treatment | 0.0687 | 0.1381 | -0.5 | 0.62 |
| block (centered) | **1.3495** | **0.1767** | **7.64** | **< 0.001** |
| **Random effects** | **Var** | **SD** | | |
| Group plate | 0.00008 | 0.009 | | |

Lifespan was analysed by a survival analysis using the coxme function, with worms that matricided or were lost entered as right-censored data points. Covariates were centred as described in the main text. Effects associated with a p value smaller than 0.05 are highlighted in bold.
