## Supplementary material for "Evolutionary causes of lifespan extension by dietary restriction: linking theory and mechanisms": Table S2

**S2:** Full model summary for lifetime reproduction in offspring of N2 parents treated with RNAi *bec-1* versus controls.

**Lifetime reproduction in N2 offspring *(N = 102)*** **linear mixed effects model**

| **Fixed effects** | **Coef** | **SE (coef)** | **ddf** | **t** | **p** |
|---|---|---|---|---|---|
| Intercept | **243.169** | **10.039** | **9.744** | **24.223** | **<0.001** |
| treatment | -11.06 | 11.742 | 93.395 | -0.942 | 0.3486 |
| block (centered) | **-60.801** | **16.818** | **5.327** | **-3.615** | **0.0137** |

| **Random effects** | **Var** | **SD** |
|---|---|---|
| Development plate | 527.5 | 22.97 |
| Residual | 3049.7 | 55.22 |

Coefficients, standard errors, test statistics and variance components are taken from a LMM on the number of offspring produced. Effects associated with a p-value smaller than 0.05 are highlighted in bold.
