## Supplementary material for "Evolutionary causes of lifespan extension by dietary restriction: linking theory and mechanisms": Table S1

**S1:** Full model summary for lifespan.

| **Lifespan *(N = 679)*** | | **coxme** | | |
|---|---|---|---|---|
| **Fixed effects** | **Coef** | **SE (coef)** | **z** | **p** |
| treatment (centered) | **0.496** | **1.6415** | **4.1** | **< 0.001** |
| strain (centered) | **-0.483** | **0.1197** | **-4.03** | **< 0.001** |
| treatment:strain | 0.462 | 0.2384 | 1.94 | 0.053 |
| **Random effects** | **Var** | **SD** | | |
| Block (3 levels) | 1.055 | 1.114 | | |
| Group plate | 0.324 | 0.105 | | |

Lifespan was analysed by a survival analysis using the coxme function, with worms that matricided or were lost entered as right-censored data points. Covariates were centred as described in the main text. Effects associated with a p-value smaller than 0.05 are highlighted in bold.
